# Identification of clinical combination therapies to induce durable responses in kidney cancers

**DOI:** 10.1101/2020.07.25.221507

**Authors:** Hui-wen Lue, Daniel S. Derrick, Soumya Rao, Anna Van Gaest, Larry Cheng, Jennifer Podolak, Samantha Lawson, Changhui Xue, Devin Garg, Ralph White, Christopher W. Ryan, Justin M. Drake, Anna Ritz, Laura M. Heiser, George V. Thomas

**Author notes:** These authors contributed equally (Hui-wen Lue, Daniel S. Derrick).

## Abstract

The lack of effective treatment options for advanced non-clear cell renal cell carcinoma (NCCRCC) is a critical unmet clinical need. Applying a high throughput drug screen to multiple human kidney cancer cells, we identified the combination of the VEGFR-MET inhibitor cabozantinib and the SRC inhibitor dasatinib acted synergistically in cells to markedly reduce cell viability. Importantly, the combination was well tolerated and caused tumor regression *in vivo*. Transcriptional and phosphoproteomic profiling revealed that the combination converged to downregulate the MAPK-ERK signaling pathway, a result not predicted by single agent analysis alone. Correspondingly, the addition of a MEK inhibitor synergized with either dasatinib or cabozantinib to increase its efficacy. This study, by employing approved, clinically relevant drugs provides the rationale for the design of effective combination treatments in NCCRCC that can be rapidly translated to the clinic.

## Introduction

RCC accounts for more than 400,000 new cancer cases and 175,000 deaths per year worldwide^1-3^.Compared to other cancers, the pathology and genetics of kidney cancer is uniquely linked^4, 5^.Approximately 75% of kidney cancers are predominantly composed of clear cells and characterized by increased angiogenesis due to the loss of the *VHL* tumor suppressor gene. Consequently, clear cell RCC (CCRCC) are responsive to drugs that that directly or indirectly inhibit angiogenesis^6-8^.The remaining ∼25% of kidney cancers, while pathologically heterogeneous (e.g. papillary, chromophobe, sarcomatoid) have an intact *VHL* gene, and are broadly classified as non-clear cell RCC (NCCRCC). NCCRCC show minimal responses to antiangiogenics and have no effective treatment options^9-16^.While recent advances in immunotherapeutic approaches have further improved outcomes for metastatic CCRCC, standard therapies for advanced NCRRCC are lacking and long-term survival is poor^4^.

We and others identified SRC, an intracytoplasmic tyrosine kinase, as a novel therapeutic target in RCC^17,18^.Despite its promise, this target has shown minimal efficacy: notably, the FDA-approved SRC inhibitor dasatinib is primarily cytostatic and fails to kill RCC cells^18^; similarly, clinical activity of SRC inhibitors have been modest with rare durable responses^19-22^.This latter observation may be because SRC is activated non-mutationally through its interactions with growth factor receptors, where it acts as a rheostat for multiple signaling pathways that mediate proliferation and survival^23^.Consequently, upfront combinatorial drug therapies that blocks SRC and its key signaling partner(s) could potentially be more effective than single agent dasatinib.

Here, we took a systematic approach toward identifying co-targeting strategies for dasatinib by performing a combination drug screen using a chemogenomic library of mechanistically annotated, clinically-relevant approved and investigational drugs that inhibit pathways involved in growth, metabolism, and apoptosis in eight representative human RCC cell lines (5 *VHL* Wild Type and 3 *VHL* Null), resulting in a dataset incorporating 37,888 single agent and dasatinib combination responses. Based on this dataset, we selected 28 promising drugs to undergo further combination matrix screening covering 6,720 distinct drug-dasatinib combinations to enable the identification of potential synergies. These studies revealed cabozantinib as a promising drug combination with dasatinib. Cabozantinib inhibits several tyrosine kinases which are biologically relevant in RCC, including VEGFRs, MET, and AXL^24^.Cabozantinib is approved for use in advanced renal cell carcinoma, having demonstrated improved progression-free survival (PFS) vs. standard-of-care sunitinib as first-line treatment in patients with intermediate- or poor-risk metastatic CCRCC^25^, and showing significant improvements in PFS, objective response rate (ORR) and overall survival (OS) when compared with everolimus in patients treated with prior anti-angiogenic therapy^26,27^.Subsequently, we performed *in vivo* testing in representative *VHL* WT RCC models. Strikingly, while single agent dasatinib and cabozantinib recapitulated the clinical responses to restrain tumor growth, the combination caused marked tumor regression. Comprehensive integration of transcriptome and phosphoproteomic analysis of the combination therapy revealed rewiring of the kinome, with inhibition of MAPK signaling required for cytotoxic synergy. Taken together, our studies have identified promising drug combinations that transcend lineage and genetic landscape to induce cytotoxicity, suggesting broad utility across different kidney cancer subgroups.

## Results

### Comprehensive high-throughput drug synergy screen

Hypothesizing that the purely cytostatic response observed with SRC inhibition alone necessitates co-targeting of bypass signaling pathways, we performed a combination drug screen to identify drugs that synergized with dasatinib to kill cancer cells **(Figure 1A; Table S1).** First, we screened a library of 296 structurally diverse, medicinally active, and cell permeable small molecules (including inhibitors of key cancer-relevant targets, e.g., VEGFR, MET, EGFR, PDGFR, PI3K, CDK and apoptosis inducing molecules, e.g. BCL2, TP53, MDM2, survivin) in eight human RCC cell lines (*VHL* WT: ACHN, SN12C, TK10, UO31, CAKI-1, and *VHL* Null: 786-0, A498, 769-P)^28^.We tested 8 doses of each drug with or without dasatinib and read viability after 5 days of drug treatment, thereby generating 37,888 single agent and drug+dasatinib dose responses. Eighty-one of the drugs passed the “Highest Single Agent” (HSA) filter, where the combination has at least 10% greater inhibition than either dasatinib or the single agent alone at the same dose, for at least three doses^29^.The viability readings were used to calculate the GI50 (the drug concentration necessary to inhibit growth by 50% compared to the untreated condition), minimum viability, and the AUC (area under the dose-response curve) between drug alone and drug+dasatinib. Next, we determined the leads for secondary screening with the following rationale: a drug should have an effect in multiple cell lines, but an effect across all cell lines is not necessary. Therefore, for a specific measurement such as GI50 or AUC, we considered drugs that were in the top 50% of the measurement for all drugs in more than half of the cell lines (i.e. 5 or more cell lines), and drugs that were in the top 25% of the measurement for all drugs in more than one cell line (i.e. 2 or more cell lines). We applied these criteria to three calculations (GI50 fold change, AUC difference between drug alone and drug+dasatinib, and AUC percent change between drug alone and drug+dasatinib) and noted the drugs that passed the criteria for multiple measurements. We shortlisted the drugs based on the above screen criteria for efficacy, safety considerations and clinical utility, and nominated 28 drugs. Notably, these drugs were active against targets relevant to RCC biology including CDK, mTOR, PI3K, MET and VEGFR. In a composite analysis of all cell lines, cabozantinib emerged as the strongest sensitizer across all parameters **(Figure 1B, C, Figures S1-3;** *see Materials and Methods*). These 28 drugs showed strong curve shifts in combination relative to single agents (drug+dasatinib/drug alone log fold change <1; **Figure 1D-K**)

**Figure 1:**
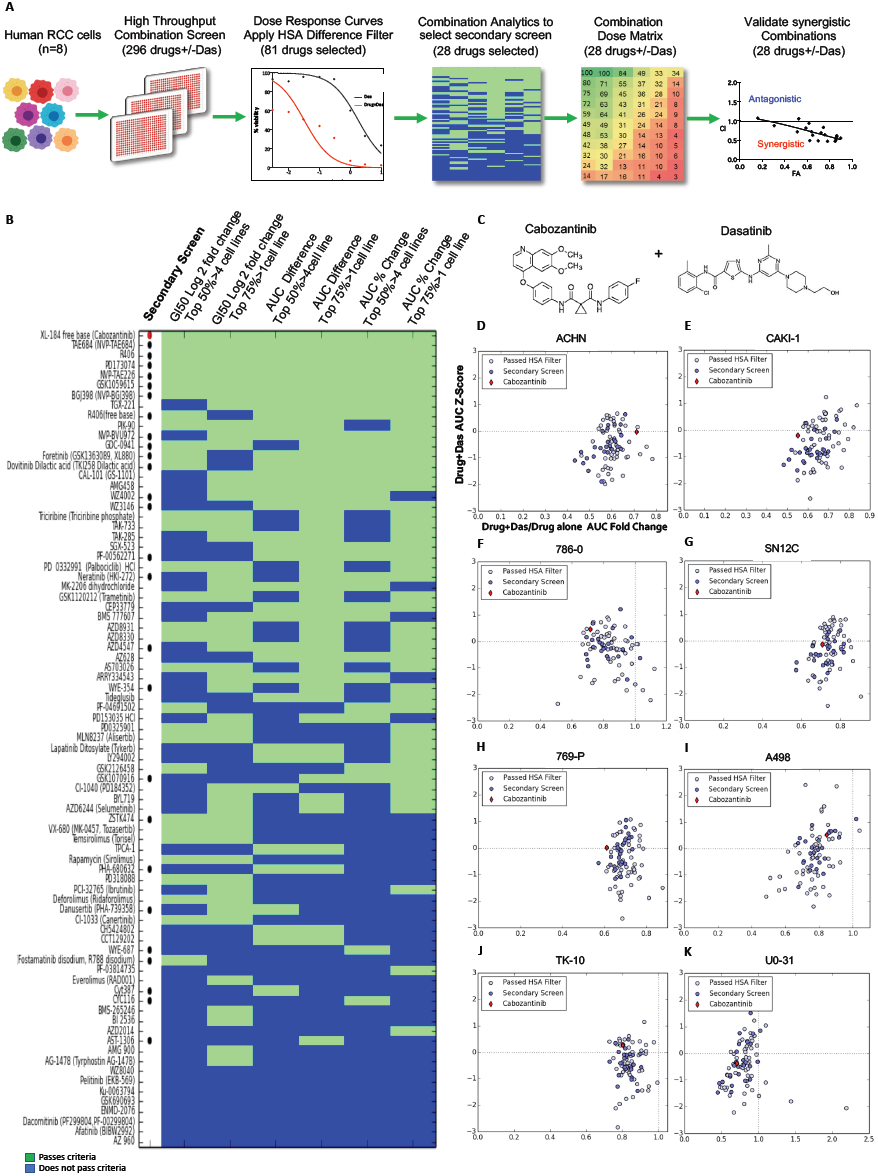
High throughput drug combination drug screen to identify sensitizers to SRC inhibition in human kidney cancer cells. **A.** Schematic of the screen workflow: Details of the primary screen with 296 drugs +/- dasatinib in 8 cell lines are in Figure S1 and Table S1. See Results for additional details. **B.** Heat map of the combination screen of 81 drugs depicting the relative sensitivity of human kidney cancer cells. The drugs shown passed the “Highest Single Agent” (HSA) filter, where the combination needs to have at least 10% greater inhibition than either dasatinib or the drug alone at the same dose, for at least three doses. Each row depicts a drug’s response according to three different measurements: G150 fold change (Columns 1 and 2), AUC difference between drug alone and drug+dasatinib (Columns 3 and 4), and AUC percent change between drug alone and drug+dasatinib (Columns 5 and 6). For each criterion, drugs pass (green) or do not pass (blue) two thresholds. In the first threshold, the drug’s measurement appears in the top 50% of all measurements in more than 4 cell lines (odd columns). In the second threshold, the drug measurement appears in the top 25% of all measurements in more than one cell line (even columns). Drugs selected for the secondary screen are denoted with a dot, with cabozantinib labeled with a red dot. **C.** Chemical structure of cabozantinib and dasatinib. **D-K.** Scatter plots denote the fold change of drug+dasatinib AUC to drug alone AUC (x-axis) versus the drug+dasatinib AUC Z-score (y-axis) for every drug. Red diamond indicates cabozantinib; dark blue dots indicate drugs selected for the secondary screen; remaining dots indicate the drugs that pass the HSA filter but were not in the secondary screen. Horizontal dashed line indicates the mean AUC Z-score of 0, and vertical dashed line indicates the AUC fold change of 1 (denoting that the drug+dasatinib AUC and the drug alone AUC are the same); **D:** ACHN, **E:** CAKI-1, **F:** 786-0, **G:** SN12C, **H:** 769-P, **I:** A498, **J:** TK10 and **K:** UO-31.

To confirm the findings of the high throughput screen, we generated dose response curves for cabozantinib alone and with the IC25 (quarter maximal inhibitory concentration) of dasatinib and observed a leftward shift with corresponding decreases in cell viability. **(Figure 2A-D)**. Next, we treated RCC cells with increasing doses of cabozantinib and dasatinib alone and in combination and observed a synergistic interaction in suppressing proliferation **(Figure 2E-H)**. Subsequently, the 28 selected drugs were further subjected to the secondary screening, which involved a dose-matrix of 6×8 in five human RCC cell lines (ACHN, CAKI-1, SN12C, 786-0 and 769-P), generating a further 6,720 dose-response signatures **(Figure S4)**. The growth inhibition values from this secondary screen across different drug doses and combinations were first analyzed for synergy using the Bliss independence model^29^.Positive Bliss scores indicate combination effects where the effect is greater than additive. Cabozantinib was identified as one the most synergistic combinations with dasatinib **(Figure 2I)**. We additionally analyzed for synergy using the multiple drug dose-median effect model as described by Chou and Talalay (CalcuSyn 2.0, Biosoft)^30^.Calcusyn calculates Combination Index (CI) for drug combinations: CI < 1 is synergistic; CI = 1 additive and CI > 1 is antagonistic **(Figure 2J, K)**. We calculated CI for 28 drug combinations in five cell lines and ranked the drugs based on their synergistic effects in combination with dasatinib. This demonstrated consistent synergy (CI<1) between dasatinib and the top ranked drugs from the primary screen, validating our selection criteria. In particular, we observed synergy between dasatinib and inhibitors of VEGFR (cabozantinib, PD1703074, foretinib, R406-fostamatinib) and PI3K (GSK1059615, and GDC-0941). In agreement with the Bliss model, cabozantinib was one of the highest ranked synergistic combinations. Notably, we confirmed our prior finding of synergy between SRC and STAT3 inhibition (CYT387), demonstrating the robustness of our screen^31^.Collectively, these experiments suggested several potential therapeutic partners for dasatinib. We selected cabozantinib for further evaluation because it has been approved for both frontline and second-line treatment in CCRCC^25-27^,is being actively studied in ongoing single and combination clinical trials (NCT03541902, NCT04022343, NCT03937219, NCT03635892), and its distinction as consistently being a top hit throughout our screens in all cell lines tested.

**Figure 2.**
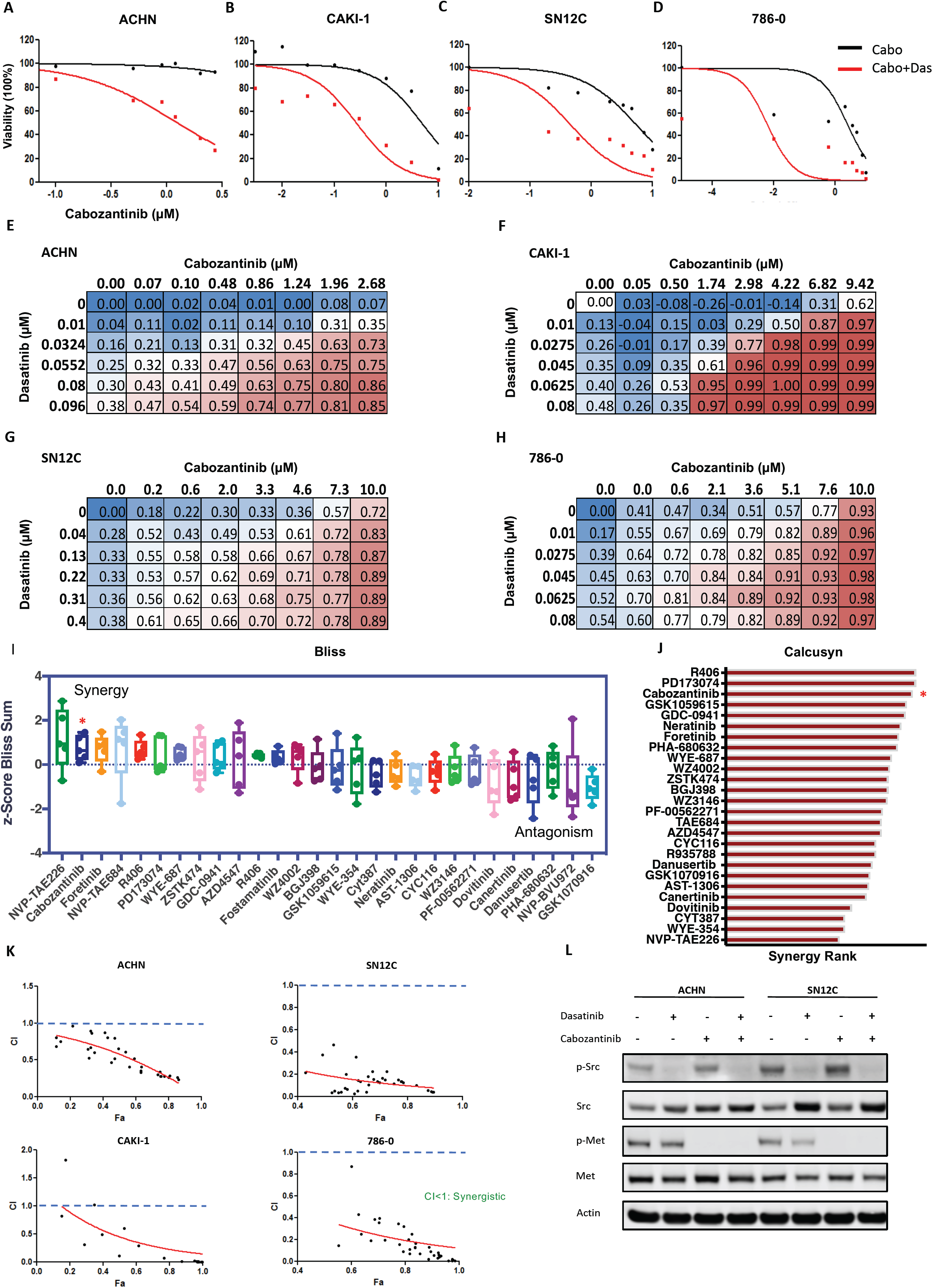
Validation of cabozantinib-dasatinib combination across representative RCC cells. **A-D.** Cell viability was assessed by Cell Titer-Glo in **A:** ACHN; **B:** CAKI-1; **C:** SN12C and, **D:** 786-0 human kidney cancer cells treated with escalating doses of cabozantinib alone (black line) or cabozantinib and a fixed dose of dasatinib at its IC_25_: 20nM for ACHN, CAKI-1, 786-0; 100nM for SN12C (red line) and dose-response curves were generated. Data represent the mean of three replicates. **E-H.** Secondary screening dose matrix of cabozantinib and dasatinib in E: ACHN; F: CAKI-1; G: SN12C and, H: 786-0 human kidney cancer. Viability was assessed after 4 days. Percent inhibition at each dose of the drug is presented. **I-K.** Dose matrices for five human RCC cell lines, ACHN, CAKI-1, SN12C, 786-0 and 769-P were generated in a 6 x8 format (6 doses of dasatinib and 8 doses of the drug) and assessed for viability after 4 days of treatment, and subjected to the estimation of synergy using the Bliss Independence Model and Calcusyn. **I.** Positive Bliss scores indicate combination effects where the effect is greater than additive. **J.** Calcusyn calculates Combination Index (CI) for drug combinations: CI < 1 is synergistic; CI = 1 additive and CI > 1 is antagonistic effects. The response of each cell line to the combination was analyzed for synergy and ranked by number of combinations that were synergistic (see Methods). **K:** CI for the cabozantinib+dasatinib combination in ACHN, CAKI-1, SN12C and 786-0 cells. **I.** ACHN and SN12C human kidney cancer cells were seeded and treated with either dasatinib (50nM) or cabozantinib (10uM), either alone or in combination. Lysates were made after 24 hr of treatment and probed with the indicated antibodies.

Because our screening data was performed using CellTiter-Glo, which is adenosine triphosphate-dependent, we additionally examined the dasatinib-cabozantinib combination using alternative assays measuring caspase activation and resazurin metabolism (CellTiter-Blue). In agreement with the CellTiter-Glo data, combined treatment with dasatinib and cabozantinib caused a marked increase in apoptosis and decrease in proliferation in RCC cells in the panel compared to dasatinib alone **(Figure S5).** Western blot analysis of human NCCRCC ACHN and SN12C cells treated with cabozantinib and dasatinib confirmed the inhibition of their respective targets, as indicated by the de-phosphorylation of MET and SRC **(Figure 2L).**

### Dasatinib-cabozantinib co-treatment induces tumor regression in human NCCRCC xenograft models

We next examined the safety and efficacy of dasatinib and cabozantinib cotreatment *in vivo* in two xenograft tumor models. While dasatinib and cabozantinib exhibited anti-tumor effect on ACHN and CAKI-1 human RCC xenografts, the combination potently inhibited tumor growth and caused tumor regression **(Figure 3A, B:** ACHN xenograft tumors**; Figure 3E, F:** CAKI-1 xenograft tumors). Importantly, the combination was well tolerated with no weight loss recorded **(Figure S6)**. Consistent with our prior reports and current *in vitro* findings, dasatinib alone had a minimal impact on apoptosis^18, 31^.In marked contrast, combination treatment with dasatinib and cabozantinib resulted in a significant increase in the magnitude of apoptosis (established by an increase in cleaved caspase 3; *P* < 0.0001) [ACHN xenograft tumors], [CAKI-1 xenograft tumors]) and a reduction in proliferation (demonstrated by a decrease in Ki-67; *P* < 0.0001) **(Figure 3C, D, I:** ACHN xenograft tumors**; Figure 3G, H:** CAKI-1 xenograft tumors). Pharmacodynamic studies demonstrated that combination therapy led to the suppression of SRC and Met-phosphorylation in treated NCCRCC xenograft tumors **(Figure 3J)**. These data support the combination of dasatinib and cabozantinib as an effective strategy for NCCRCC.

**Figure 3.**
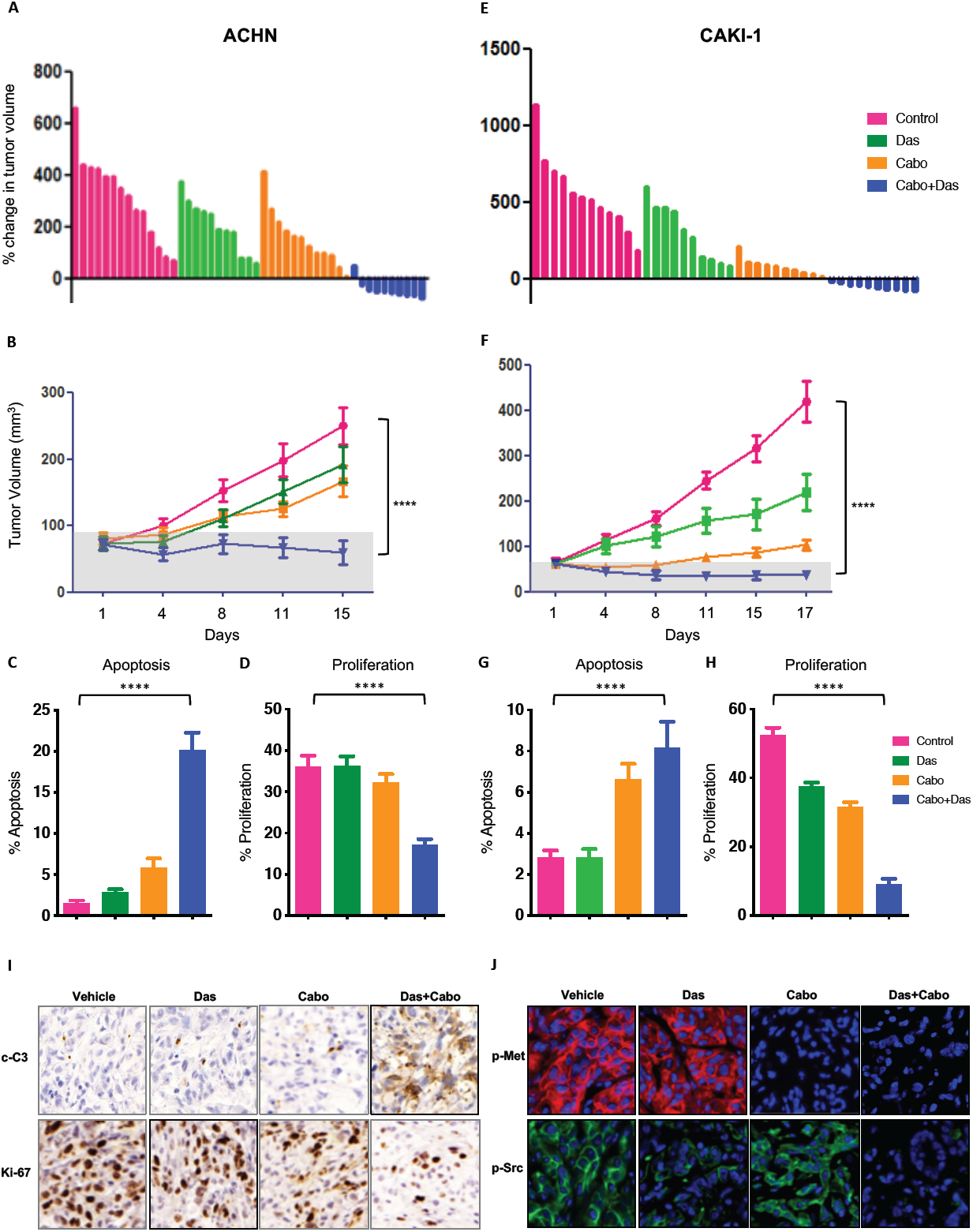
Cabozantinib combines with dasatinib to induce tumor regression. **A-D.** ACHN xenografts treated with vehicle, dasatinib (25mg/kg/day), cabozantinib (30mg/kg/day) and dasatinib-cabozantinib (25mg/kg/day+30mg/kg/day) combination. **A:** Waterfall representation of response of each tumor after 15 days of treatment is shown. **B:** Tumor volume is shown. Error bars represent mean ± SEM; (n>8 per treatment group), shaded area represents regression; (control vs cabozantinib+dasatinib P<0.0001****) **C:** Effect on apoptosis (c-C3) and, **D:** proliferation (KI67) in ACHN xenograft tumors. Error bars represent mean ± SEM. (**C:** control vs cabozantinib+dasatinib, p<0.0001****; **D:** control vs cabozantinib+dasatinib, p<0.0001****). **E-H.** CAKI-1 xenografts treated with vehicle, dasatinib (35mg/kg/day), cabozantinib (10mg/kg/day) and dasatinib-cabozantinib (35mg/kg+10mg/kg/day) combination. **E:** Waterfall representation of response of each tumor after 15 days of treatment is shown. **F:** Tumor volume is shown. Error bars represent mean ± SEM; (n>8 per treatment group), shaded area represents regression; (control vs cabozantinib+dasatinib P<0.0001****) **G:** Effect on apoptosis (c-C3) and, **H:** proliferation (KI67) in CAKI-1 xenograft tumors. Error bars represent mean ± SEM. (**G:** control vs cabozantinib+dasatinib, p=0.0002****; **H:** control vs cabozantinib+dasatinib, p<0.0001****). **I.** Representative images of tumor tissue from ACHN xenografts treated with the indicated drug regimens were evaluated by immunohistochemistry for cleaved-Caspase 3 and KI-67. **J.** Representative images of tumor tissue from ACHN xenografts treated with the indicated drug regimens were evaluated by immunofluorescence for p-MET and p-SRC.

### Combination treatment of NCCRCC cells reveals deterministic and stochastic signaling outputs

To obtain further insight into the signaling pathways acutely affected by dasatinib and cabozantinib co-treatment, we evaluated the changes in the phosphoproteome of the human NCCRCC cell line ACHN after dasatinib, cabozantinib and the combination treatment via quantitative, label-free quantitative mass spectrometry^32-34^.Supervised hierarchical clustering revealed duplicate samples clustered together, but that treatment altered phosphorylation levels of phosphopeptides with 3,369 phosphoserine and phosphothreonine (pST) peptides and 81 phosphotyrosine (pY) peptides significantly differed between treated and untreated cells (FDR <0.10, 0.20, respectively) **(Figure 4A)**. As expected based on known dasatinib and cabozantinib targets, the SRC family kinases (including SRC, Fyn, Hck, Lyn, and Fgr) and KDR (VEGF) were predicted to be significantly less active in the combination-treated cells based on pY kinase substrate enrichment analyses (KSEA)^33,35^,an approach that estimates changes in a kinase’s activity based on the collective phosphorylation changes of its identified substrates. Focusing on the single agents, cabozantinib alone enriches for phosphatase activity of PTPN2/6, tyrosine kinases activity of RET and ERBB3/4, and transcription factor activity of STAT1/3. Dasatinib treatment alone also enriches phosphatase activity of PTPN6 and FLT1 kinase activity, a binding protein to VEGFR for activation. The robustness of the data and analysis was demonstrated by the de-enrichment of Abl1 activity, a known target of dasatinib^36^.These changes in kinase/phosphatase activity enrichment suggest for potential resistance mechanisms for cell survival when given either single agent **(Figure 4 B, C: left panels)**.

**Figure 4:**
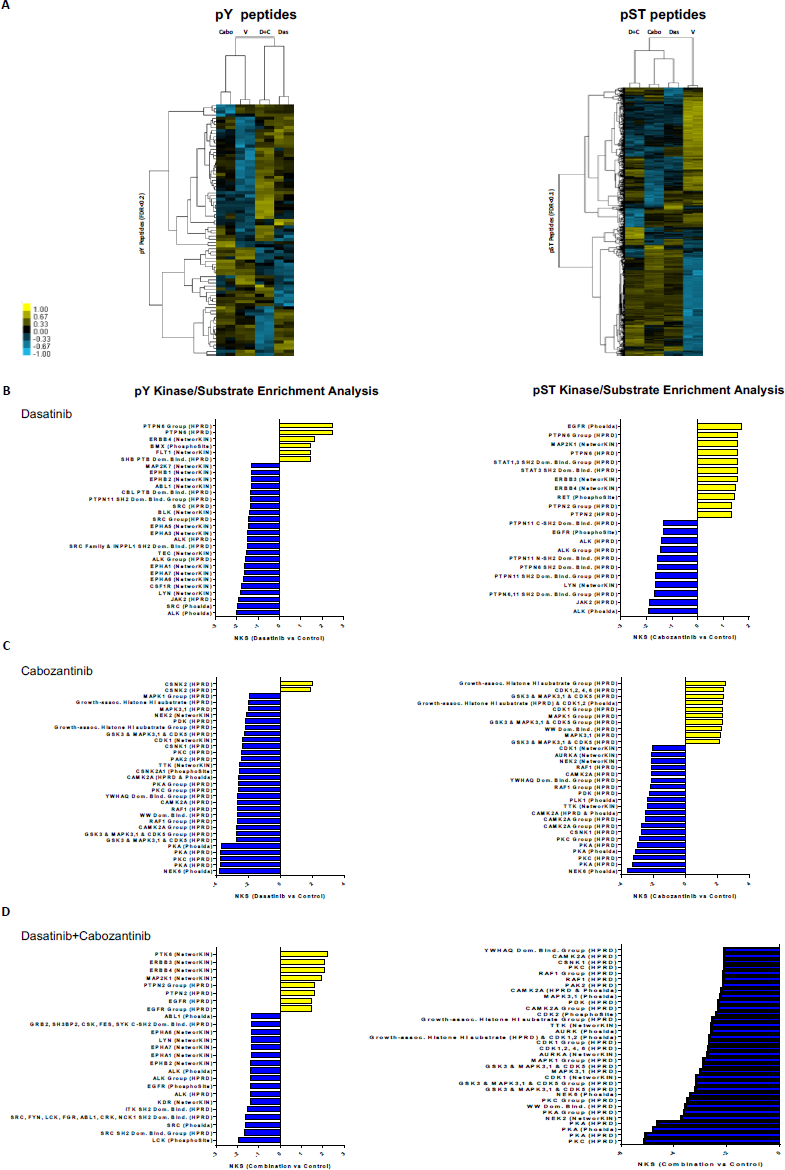
Characterization of the phosphoproteome in cabozantinib-dasatinib co-treated NCCRCC cells. **A.** Supervised hierarchical clustering heatmaps of: phosphotyrosine peptides (pY, left panel), and phosphoserine and phosphothreonine peptides (pST, right) and identified from cabozantinib, dasatinib and the combination in treated and untreated ACHN human RCC cells with two technical replicates. 81 unique pY phosphopeptides (rows) and 3,369 unique pST phosphopeptides were either 4-fold more enriched or 4-fold less enriched, on average (pY: FDR <0.2; pST: FDR<0.1; t-test p<0.2), in combination-treated cells compared to untreated cells. **B-D.** Kinase-substrate enrichment analysis (KSEA) of **B:** dasatinib; **C:** cabozantinib; **D:** cabozantinib-dasatinib co-treated and untreated pY (Hits ≥ 3; FDR <0.05; left panels). and pST data (Hits ≥ 30; FDR <0.01; right panels). Positive NKS (Normalized Kolmogorov-Smirnov Score) infer greater kinase activity in cabozantinib-dasatinib co-treated cells while negative NKS indicate greater activity in untreated cells (Unfiltered summary are in **Tables S2-S7**)

KSEA for the pST motifs for the dasatinib-cabozantinib combination showed striking decreased activity across multiple kinases, including CDKs, NEK2, PKA and PKCs that would not have been predicted from the known kinome single activity profile of dasatinib or cabozantinib. However, it has been shown in previous literature that CDKs are phosphorylated and activated by SRC kinase, increasing the entry to cell cycle via phosphorylation and degradation of p27. In addition, BCR-ABL can also function to activate CDKs via p27 as well. Both of which is inhibited with dasatinib^37, 38^.Conversely, we observed enrichment of PTK6, ERBB, MAP2K1 pY motifs, all of which may interact with one another, suggesting potential activation of bypass tracks from combination therapy **(Figure 4 D: right panels)**. Collectively, the phosphoproteome data provide strong evidence that the combination treatment, in addition to reducing known SRC and KDR signaling, also impacts multiple unexpected signaling networks that may be direct or indirect targets of these drugs.

### Integration of transcriptomic and phosphoproteomic datasets reveals coordinated inhibition of MAPK-ERK pathway

Next, we sought mechanistic understanding for the observed tumor regressions after dasatinib-cabozantinib co-treatment by performing RNA sequencing of ACHN human NCCRCC cells treated with dasatinib, cabozantinib and the combination for 24 hours. Cabozantinib treatment caused differential expression of 4,026 (2,248 up + 1,178 down) genes compared to vehicle treatment. Dasatinib treatment resulted in a much more modest transcriptional response relative to cabozantinib treatment, inducing differential expression of 49 (48 up + 1 down) genes compared to vehicle treatment. Co-treatment induced the greatest transcriptional response, resulting in differential expression of 5,839 (3,048 + 2,791) genes compared to vehicle (Log2FC ≥ 0.5 or ≤ 0.5; q < 0.01), 65% of which were shared with either or both of the individual treatments **(Figure 5A)**. Comparison of overall expression profiles across the treatments revealed a high degree of correlation across treatments (**Figure S7**; *Methods*). These results indicate shared molecular programs between each treatment, but also point to molecular changes specifically induced by the combination treatment.

**Figure 5:**
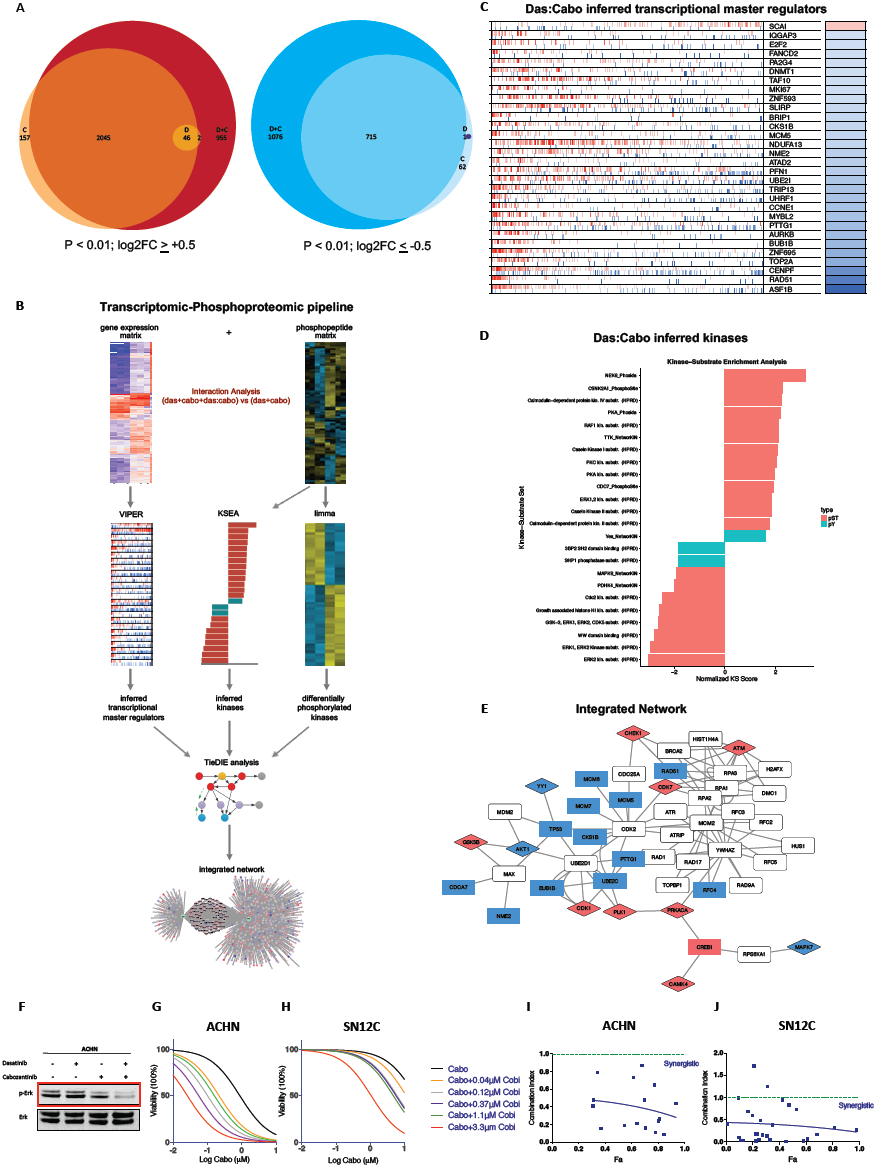
Cabozantinib and dasatinib converge to downregulate the MAPK-ERK signaling pathway. **A.** ACHN cells were treated with 50nM cabozantinib, 10uM dasatinib or the combination for 24hrs. Differentially expressed genes induced by single or combination drug treatment. Euler diagrams show overlaps in genes with significant increase (log2FC ≥ 0.5; FDR-corrected p-value ≤ 0.01) or decrease (log2FC ≤-0.5; FDR-corrected p-value ≤ 0.01) in expression following individual or combination drug treatment. **B.** Transcriptomic-Phosphoproteomic Data Integration Workflow: To identify genes and phosphopeptides selectively affected by the cabozantinib-dasatinib drug interaction, we compared full models, including terms for the individual drugs and their interaction, to reduced models that only model the individual drug effects. Genes and phosphopeptides were then ranked by the extent to which their expression was better explained by inclusion of an interaction term. Using these ranked lists, we used the VIPER algorithm to infer Master Regulator activity from transcriptional profiling data, and we used the KSEA algorithm to infer upstream kinase activity from the phosphoproteomic data. The TieDie algorithm was used with the Multinet interaction network to combine inferred transcriptional master regulators, inferred kinases, and directly measured kinases into an integrated network associated with response to the combination drug treatment. **C.** Inferred master regulators induced by cabozantinib-dasatinib combination: Transcriptional master regulators driving the unique response to combination treatment were inferred from genes ranked by the interaction coefficient using VIPER. Hashmarks in each row represent the positions of the regulon genes in a list of all genes ranked by the interaction coefficient. Red marks indicate positive targets; blue marks indicate negative targets. Heatmap on the right indicates, for each Master Regulator, the direction of enrichment (blue = negative; red = positive). The top 30 significant Master Regulators are shown. **D.** Inferred kinases induced by cabozantinib-dasatinib treatment: Kinase activity driving the effect of drug interaction was inferred from the phosphoproteomic data ranked by the interaction coefficient using Kinase-Substrate Enrichment Analysis (KSEA) (q-value < 0.05). **E.** Integrated Transcriptomic/Phosphoproteomic cabozantinib-dasatinib Interaction Network: Cabozantinib-dasatinib interaction network generated from VIPER master regulator enrichment scores, KSEA kinase enrichment scores, and kinase using Tie-Die algorithm and MultiNet interaction network. Red nodes have increased expression due to interaction effect; blue nodes have decreased expression due to interaction effect. Rectangles indicate transcription factors and diamonds indicate kinases. **F.** ACHN human kidney cancer cells were seeded and treated with either dasatinib (50nM) or cabozantinib (10uM), either alone or in combination. Lysates were made after 24 hr of treatment and probed with the indicated antibodies. **G, H.** Dose response curves of cell viability of human NCCRCC cell lines, ACHN, SN12C to varying doses of cabozantinib and cobimetinib after 72hrs exposure. **I, J.** Calculated median effect drug synergy CI scores (Calcusyn) across cabozantinib and dasatinib combinations. Horizontal dashed line indicates a CI=1, where points below the line indicate synergy and points above indicate antagonism

To hone in on the genes and phosphopeptides changes specifically induced by the combination treatment, we developed a statistical and network approach to compare across these treatments **(Figure 5B)**. We used likelihood ratio tests to compare “full” linear models including terms for effects of dasatinib, cabozantinib, and their interaction to “reduced” models, which only included terms for cabozantinib and dasatinib individually. We prioritized genes and phosphopeptides that were significantly better explained by the full model. Three hundred and twenty-four genes were significantly better explained after the inclusion of an interaction term, indicating that these genes are preferentially activated after combination treatment (likelihood ratio test: q < 0.01). To identify molecular nodes that may mediate the response to the combination treatment, we performed a master regulator (MR) analysis of the 324 genes to identify transcription factors or signaling molecules that serve signal integration hubs. Specifically, we used the MARINa (Master Regulator Inference algorithm) and VIPER (Virtual Proteomics by Enriched Regulon analysis) algorithms and a gene regulatory network derived from the TCGA KIRP cohort to infer Master Regulator activity from genes ranked by the effect of interaction of expression^39, 40^**(Figure 5C)**. The dasatinib-cabozantinib interaction inferred transcriptional MRs were involved in DNA replication and repair, transcriptional regulation, proliferation and mitosis, e.g. ASF1B, RAD51, CENF, TOP2A, BUB1B, AURKB, PTTG1, MCM5 and CCNE1^41,42^. Accordingly, gene set enrichment analysis (GSEA) of the inferred transcriptional MRs demonstrated downregulation for cell cycle, DNA replication, homologous recombination and base excision repair^43^ **(Figure S8).**

We used a similar analytical approach to analyze the proteomic data and identified 959 proteins that could be explained by an interaction between dasatinib and cabozantinib. Kinase activity driving the effect of drug interaction was inferred from the phosphoproteomic data using kinase-substrate enrichment analysis (q-value < 0.01), and demonstrated concordant downregulation of cell cycle associated kinases CDK1/CCD2. Notably, KSEA also suggested downregulation of multiple nodes of the MAPK pathway, e.g. SHP, ERK1, ERK2, ERK5 (MAPK7), JNK1 (MAPK8) with the dasatinib-cabozantinib combination^44^ **(Figure 5D)**.

To integrate across transcriptional and phosphoproteomic datasets, we leveraged the TieDIE algorithm with the Multinet interaction network to combine inferred transcriptional MRs, inferred kinases, and kinases into an integrated network associated with response to the combination drug treatment^45^. TieDIE analysis predicted a complex regulatory network underlying the response to the cabozantinib-dasatinib combination treatment that converged to downregulate the MAPK signaling pathway. Specifically, it connected downstream nuclear translocating proteins of the MAPK pathway such as MAPK7 (ERK5), RPS6KA1 (P90-RSK1) to the inferred transcriptional MRs involved in DNA replication and proliferation (e.g., BUB1B, MCM5, PTTG1; **Figure 5E**)^46^.Critically, in support of the TieDIE “interactome”, western blot analysis of ACHN cells treated with dasatinib or cabozantinib singly demonstrated only minimal effect on MAPK activity (as determined by ERK1/2 phosphorylation), while the combination markedly suppressed MAPK activity **(Figure 5F)**.

To better define the effect of MAPK pathway inhibition, we evaluated the addition of MEK inhibitors to either dasatinib or cabozantinib. First, we observed that trametinib (an approved MEK inhibitor) when combined with dasatinib induced substantial decreases in proliferation in multiple kidney cancer cells, as demonstrated by the shift in GI50 **(Figure S9)**. Since clinical trials testing another MEK inhibitor is already underway for kidney cancer (cobimetinib^47^: NCT NCT03264066), and with cabozantinib already approved in kidney cancer, we next evaluated the combination of cobimetinib with cabozantinib in ACHN and SN12C human NCCRCC cells. We used a dose matrix to sample a large range of concentrations and concentration ratios and saw that cobimetinib-cabozantinib combination exhibited marked decreases in cell viability at clinically relevant doses (peak plasma levels of 0.51μM and 4.61 μM respectively)^48^ **(Figure 5G, H)**. We used the Calcusyn analysis to test for synergy. This demonstrated consistent synergy [combination index (CI) < 1] between cobimetinib and cabozantinib **(Figure 5 I, J)**. Accordingly, these findings demonstrate that inhibition of the MAPK pathway contributes to the synergistic effect of the dasatinib-cabozantinib combination in NCRCC, and highlights the potential for additional novel co-targeting strategies.

## Discussion

NCCRCC accounts for ∼20% of kidney cancer, with papillary, chromophobe and sarcomatoid variants accounting for the majority of subtypes^5^.For advanced NCCRCC, the only treatment options, which have been largely extrapolated from agents studied in CCRCC, include antiangiogenics or mTOR inhibitors. However, several large studies have demonstrated that patients with NCCRCC have a worse prognosis with lower response rates to these therapies^9-16^. Consequently, the NCCN recommends enrolment to a clinical trial for NCCRCC patients as the preferred choice^49^.Taken together, the lack of treatment options for NCCRCC patients represents a critical unmet clinical need. Here, we sought to identify preclinical synergistic drug combinations that would transcend lineage and genetic landscape to induce cytotoxicity in NCCRCC and ultimately lead to deep and durable responses in patients.

We demonstrate that the dasatinib-cabozantinib combination has potent synergy in 2D cell culture models. Importantly, this combination induces tumor regression *in vivo*, confirming the robustness of our experimental screening approach. We observed that multiple NCCRCC cells appear to respond similarly to the combination, suggesting broad utility across this histologically diverse group of cancers. Mechanistically, the dasatinib-cabozantinib combination converge to suppress MAPK signaling to induce cytotoxicity. Critically, this suggests that combining dasatinib or cabozantinib with MEK inhibitors would be a viable combination. In support of this, we found that the cabozantinib-cobimetinib combination synergized to markedly reduce cell viability. Taken together, our preclinical findings provide rationale for clinical trial design.

What then is the most straightforward path to the clinic? We speculate that these findings have direct clinical implications for cabozantinib, providing rationale for further clinical study of cabozantinib in NCCRC and supporting future combination treatments. Retrospective studies suggest that cabozantinib has activity in patients with papillary RCC (the commonest NCCRCC subtype, accounting for ∼15% of all kidney cancers), has activity in metastatic RCC to bone^50^, and prospective clinical studies are ongoing (NCT02761057). However, it is highly likely that NCCRCC patients will develop resistance to single agent cabozantinib, either due to short term signaling adaptations (e.g. bypass tracks) or longer-term selection of resistance variants (e.g. gatekeeper mutations). Combination studies with cabozantinib need to be considered, and our data suggest that either dasatinib or cobimetinib would be suitable. Additionally, cobimetinib is currently undergoing combination studies in RCC (+ Azetolizumab, NCT03264066). Importantly, our work provides the mechanistic rationale on how best to develop these combinations for the clinic.

The caveats here are limitations of cell culture and mouse xenograft studies, and how these can be extrapolated to clinical trials. While the dasatinib-cabozantinib combination was well tolerated in mice, the potential clinical toxicities in the setting of patients who have exhausted multiple lines of therapies and with co-morbidities are part of the real-world study considerations. Appropriate dose-finding studies are needed as a first step to explore safety and feasibility, either stand-alone or incorporated into larger efficacy trials.

## Material and Methods

### Cell lines

ACHN, A498, 769-P, 786-O, CAKI-1, SN12C, TK10 and UO31 were used in this study and were obtained from the NCI and ATCC. Cell lines were maintained in Dulbecco’s modified Eagle’s medium (DMEM) supplemented with 10% fetal bovine serum at 37°C in a 5% CO_2_ incubator.

### High throughput drug screening and workflow on the identification of hits

We did screening using a library of 296 small molecules (that included kinase inhibitors and apoptosis inducing molecules; **Table S1**) in 8 cell lines (ACHN, A498, 769-P, 786-O, CAKI-1, SN12C, TK10 and UO31). We did a 8 dilutions of drug with or without dasatinib and read viability using cell titer glo after 5 days of drug treatment. The readings were used to calculate the GI50 (using XLfit), minimum viability, AUC (area under curve), AUC difference between drug alone and drug+das and Z score values. We applied multiple criteria to identify the hits from the primary screening that were additionally validated in the secondary screening. The following are the criteria used: **1) GI50 fold change:** Drugs that have at least 5-fold change in GI50 ratio between drug alone and drug+dasatinib (ratio GI50 drug/ GI50 combination) in 50% of cell lines (n = 38). **2) AUC difference:** Drugs that have AUC difference of more than 0.9 (average is 0.9) between drug alone and drug+dasatinib in at least 50% of cell lines (n = 31). **3) AUC % change:** Drugs that have AUC % change of more than 25% (average is 25%) between drug alone and drug+dasatinib in 66% (4 of 6) cell lines (n = 31). **4) GI50 of combination:** Combinations that are in top 50% (based on GI50 z-score calculation) in 50% (3 of 6) cell lines and pass through the minimum viability cutoff of 15% across cell lines that (average min viability value is looked at) (n=18). **5) AUC of combination:** Combinations that are in the top 50% (based on AUC z-score calculation) and in 50% (3 of 6) cell lines and pass through the minimum viability cutoff of 15% across cell lines that (average min viability value is looked at) (n = 13). We shortlisted the drugs based on the above criteria and picked 28 drugs that were identified by at least 4 of the 5 parameters. These 28 drugs were further subjected to the secondary screening, which involved a dose-matrix of 6×8 (6 doses of dasatinib and 8 doses of the drug). The growth inhibition values from the secondary screening were subjected to the estimation of synergy using Calcusyn. Calcusyn calculates Combination Index **(CI)** for drug combinations: CI < 1 is synergistic; CI = 1 additive and CI > 1 is antagonistic effects^30^.We calculated CI for 28 drug combinations and ranked the drugs based on its synergistic effect in combination with dasatinib from highest to lowest.

### Cell viability and apoptosis analysis

Cell viability assays were performed by plating cells/well in 96-well plates in triplicate and treating the following day with the indicated agent: dasatinib (dose range of 0–400 nM) and cabozantinib (dose range of 0–10 μM). The experiment was continued for 3 days and then the cells were treated with CellTiter-Blue (Promega) and incubated for 1 hour. Fluorescence was measured and quantified and photographs were obtained using a Cytation 5 Cell Imaging Reader. The effect of dasatinib, cabozantinib and the dasatinib+cabozantinib combination on cell number was assessed as fold of DMSO-treated control cells. Experimental results are the average of at least three independent experiments. Apoptosis was determined using Caspase 3/7-Glo assay kit (Promega) following the manufacturer’s instructions. Briefly, 4000 cells per well were plated in 96 well plates and cultured for 24h. Cells were treated with dasatinib, cabozantinib and the combination of dasatinib+cabozantinib for 72h, and then 100 μl reagents were added to each well and incubated for 30 min at room temperature. Caspase 3/7 activity was measured using a luminometer. Luminescence values were normalized by cell numbers. The effect of dasatinib, cabozantinib and the combination of dasatinib+cabozantinib on caspase 3/7 activation was assessed as fold of DMSO-treated control cells.

### Western Blotting

Cells were plated in 6 well dishes and treated the following day with the indicated agents. Treatments were for 24 hours, after which cells were washed with ice cold PBS and lysed with RIPA buffer (Sigma). Phosphatase inhibitor cocktail set II and protease inhibitor cocktail set III (EMD Millipore) were added at the time of lysis. Lysates were centrifuged at 15,000g x 10 min at 4 degrees C. Protein concentrations were calculated based on a BCA assay (Thermo Scientific) generated standard curve. Proteins were resolved using the NuPAGE Novex Mini Gel system on 4% to 12% Bis-Tris Gels (Invitrogen). For western blotting, equal amounts of cell lysates (15-20 μg of protein) were resolved with SDS-PAGE, and transferred to membranes. The membrane was probed with primary antibodies, washed, and then incubated with corresponding fluorescent secondary antibodies and washed. The fluorescent signal was captured using LI-COR (Lincoln, NE) Odyssey Imaging System, and fluorescent intensity was quantified using the Odyssey software where indicated. The following antibodies were used for Western blots: p-SRC (Y416), SRC, p-MET (Y1234/1235), p-p44/42 MAPK (Erk1/2) (T202/Y204), ERK and β-actin (AC15) (Abcam). Ki67 (Dako) and cleaved caspase3 (Cell Signaling Technologies) were used for immunohistochemistry. Dasatinib, cabozantinib and cobimetinib were purchased from Selleck chemicals.

### *In Vivo* Xenograft Studies

6-week old mice were utilized for human renal cell carcinoma xenografts. For both ACHN and CAKI-1 cell lines 2×10^6^ cells were diluted in 50 µl of PBS and 50 µl of Matrigel (BD Biosciences) and were injected subcutaneously into the right and left flank of each mouse. Tumors were monitored until they reached an average size of 50-80mm3 (approximately 2 weeks), at which point treatments were begun. dasatinib (25mg/kg/day: ACHN; 35mg/kg/day: CAKI-1) was administered by oral gavage 5 days/week. Cabozantinib (30mg/kg/day: ACHN; 10mg/kg/day: CAKI-1) were administered by oral gavage 3 days/week. Dasatinib and cabozantinib were dissolved in NMP/PEG. Tumors and mouse weights were measured twice weekly. At least 6-8 mice per treatment group were included. All mice were euthanized using CO_2_ inhalation followed by cervical dislocation per institutional guidelines at Oregon Health and Science University. Experiments were approved by the Institutional Animal Care and Use Committee at OHSU.

### Immunohistochemistry

Immunostaining was performed following deparaffinization and rehydration of slides. Antigen retrieval was performed in a pressure cooker using citrate buffer (pH 6.0) for 4 min. Nonspecific binding was blocked using Vector mouse IgG blocking serum 30 min at room temperature. Samples were incubated at room temperature with rabbit monoclonal antibodies cleaved caspase 3 (Cell Signaling Technologies), and Ki67 (Dako). Slides were developed with Vector Immpress rabbit IgG (#MP7401) and Vector Immpress mouse IgG (Vector Laboratories) (#MP7400) for 30 min at room temperature. Chromogenic detection was performed using Vector Immpact DAB (Vector Laboratories) (#SK4105) for 3 min. Slides were counterstained with hematoxylin. A 3DHistech MIDI Scanner (Perkin Elmer) was used to capture whole slide digital images with a 20x objective. Images were converted to into MRXS files and computer graphic analysis was completed using inForm 1.4.0 Advanced Image Analysis Software (Perkin Elmer).

### Immunofluorescence

H&E slides of formalin fixed, paraffin embedded tissue was used to assess morphological integrity of tumor samples. Once integrity was confirmed, immunofluorescent analysis was performed for p-SRC (Y416), p-MET (Y1234/1235) (Cell Signaling Technologies). Four μ sections were cut, de-paraffinized and rehydrated. Antigen retrieval was performed using citrate for 4 min in a pressure cooker. Slides were blocked using 2.5% normal goat serum for 30 min then incubated in primary antibody for 1hr followed by secondary antibody mouse anti-rabbit Alexa 488 (1:1000, Molecular Probes) for 30 min. Slides were rinsed in PBS, air dried, and coverslipped using Dako mounting media with Dapi.

### RNA-Seq preprocessing and analysis

RNA-Seq libraries were prepared from triplicate 24-hour treated cell lines using the RNeasy extraction kit (Qiagen). Single-end 100bp reads were sequenced with an Illumina HiSeq 2500. Reads were trimmed of low-quality bases and adapter sequences using TrimGalore (v0.4.4) a wrapper for cutadapt (v1.10)^51^.Reads were aligned to the human reference genome (GRCh38) and summarized to gene-level abundances using STAR (v2.4.0.1)^52^.Gene expression heatmaps were created with the R package Complex Heatmap (v1.17.1)^53^.

Differentially expressed genes in drug-treated samples were identified with DESeq2 (v1.18.1)^54^ to compare each treatment (dasatinib, cabozantinib, and dasatinib + cabozantinib co-treatment) to the vehicle-treated condition. Genes with an FDR-corrected q-value ≤ 0.01 and a log2FoldChange ≥ 0.5 or ≤ -0.5 were considered differentially expressed.

Significance of the cabozantinib-dasatinib interaction effect on gene expression was assessed using a likelihood ratio test to compare two generalized linear models. Both models feature logarithmic link functions

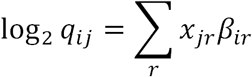

where parameter *q*_*ij*_ is proportional to the predicted true concentration of transcripts in sample *j, x*_*jr*_ are the design matrix components for sample *j*, and *β*_*ir*_ are the coefficients (log2 fold changes). The full model

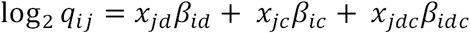

contains design matrix components and coefficients for the dasatinib effect (*x*_*jd*_*β*_*id*_), the cabozantinib effect (*x*_*jc*_*β*_*ic*_), and an interaction effect (*x*_*jdc*_*β*_*idc*_). The reduced model

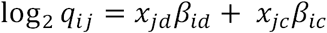

contains terms only for the dasatinib effect (*x*_*jd*_*β*_*id*_) and the cabozantinib effect (*x*_*jc*_*β*_*ic*_).

We considered genes to be affected by the interaction of the drugs if the full model explained the observed effects significantly better than the reduced model (FDR-corrected q-value < 0.01).

### Gene Set Enrichment Analysis

Enrichment of specific pathways and ontological terms in the gene most affected by the drug interaction was assessed using the Broad Gene Set Enrichment Analysis (GSEA) (v3.0) Java application^43^and the Broad MSigDB (v6.2) Hallmark gene set collection. Preranked input for GSEA was generated by ranking genes by the log2 fold change of the by the cabozantinib-dasatinib drug interaction coefficient. P-values were calculated by gene label permutation and were FDR-corrected for multiple testing. Gene sets with an FDR-corrected q-value < 0.1 were considered significant.

### Master Regulator Analysis

Master Regulator analysis was used to infer transcription factors responsible for the observed drug interaction-induced changes in gene expression. The R package VIPER (v1.12)^39^ was used to test for enrichment of Master Regulators in a list of all genes ranked by the coefficient of the interaction effect (*β*_*idc*_), using a network generated from gene expression profiles of the TCGA kidney renal papillary cell carcinoma cohort^40^. P-values were calculated by gene label permutation and were FDR-corrected for multiple testing. Master Regulators with an FDR-corrected q-value < 0.1 were considered significant.

### Phosphoproteomic drug interaction analysis

Interaction effects in the phosphoproteomic data were analyzed in R using the limma (3.36.3)^55^ package. Significance of the cabozantinib-dasatinib interaction effect on phosphoprotein levels was assessed using an empirical Bayes method to compare a full model, which models individual drug and drug interaction effects, to a reduced model, which models only the effects of the individual drugs. The drug interaction was considered to have a significant effect on phosphopeptide abundance if the FDR-corrected q-value was < 0.05.

### Data Integration using TieDie

An integrated network of inferred drug interaction effects on transcription and phosphoprotein levels was created using TieDie (v1.0)^45^ software. TieDie was used to map enriched transcription factors inferred from the RNA-seq data, kinases significantly affected by the drug interaction effect, and enriched kinases inferred from the phosphoproteomic data onto nodes of the Multinet reference network derived from protein-protein interaction, phosphorylation, metabolic, and gene regulatory networks. Inferred master regulators (FDR-corrected q-value < 0.05, NES ≥ 5) were used as the gene expression (“downstream”) input nodes. KSEA-inferred kinases (FDR-corrected q-value ≤ 0.05) were used as the kinase (“upstream”) input nodes. Additionally, directly measured kinases significantly affected by the drug interaction (FDR-corrected q-value ≤ 0.5 for pY phosphopeptides; FDR-corrected q-value ≤ 0.01 for pST phosphopeptides) were included as upstream input. A more stringent q-value threshold was used for pST phosphopeptides to reduce the number of input nodes. The input weight of inferred transcription factors and kinases was equal to their enrichment score; the input weight of directly measured kinases affected by drug interaction was equal to the interaction coefficient *β*_*idc*_. A heat diffusion algorithm was then run to model diffusion of “heat” from the input nodes to nearby nodes in the network, and a subnetwork of agreement between the data types was identified by identifying nodes that receive heat from both data sources.

### Code Availability

All analysis scripts can be found in the following GitHub repository: https://github.com/danielderrick/thomas_kidney_drug_analysis

### Phosphoproteomic screen and analysis

Phosphopeptides were enriched and analyzed by mass spectrometry as previously described in detail^32^.Briefly, cells were lysed and proteins extracted using a guanidinium-based lysis buffer. Lysates were digested using Lys-C and trypsin. Phosphotyrosine peptides were enriched via immunoprecipitation (4G10 antibody) and titanium dioxide. Phosphoserine/threonine peptides were enriched by strong cation exchange and titanium dioxide. After desalting with C18 columns, the phosphopeptide samples were subjected to liquid chromatography-tandem mass spectrometry.

### Mass spectrometry data analysis

MS raw files were analyzed via MaxQuant version 1.5.3.30 ^56^ and MS/MS fragmentation spectra were searched using Andromeda^57^ against human canonical and isoform sequences in Swiss-Prot (downloaded in August 2017 from http://uniprot.org)^58^.Quantitative phosphopeptide data were log10 transformed and missing data were imputed before applying quantile normalization as previously described^33^.Hierarchical clustering was performed on the Cluster 3.0 program^59^, using distance that is based on the Pearson correlation and applying pairwise average linkage analysis. Java Treeview was used to visualize clustering results^60^.

### Kinase substrate enrichment analysis

Kinase substrate enrichment analysis (KSEA) was performed as previously described^33, 35^.Briefly, the phosphopeptides were rank ordered by fold change, on average, between combination (dasatinib + cabozantinib) treatment vs control and the enrichment score was calculated using the Kolmogorov-Smirnov statistic. Permutation analysis was conducted to calculate statistical significance. The normalized enrichment score was calculated by dividing the enrichment score by the average of the absolute values of all enrichment scores from the permutation analysis **(Tables S2-S7)**.

### Statistical analysis

Mouse tumor size was analyzed by 2-way ANOVA with time and drug as factors, using GraphPad Prism. Mouse weight during treatment was analyzed by repeated measures 2-way ANOVA, with time and drug as factors. A *P* value less than 0.05 was considered statistically significant. Immunohistochemistry: *P*-values were calculated using one-way ANOVA, with Bonferroni’s multiple comparison test. *\* denotes P < 0.05, ** denotes P < 0.01, and *** denotes P < 0.001.* The plots here were created using R version 3.6.1. Packages needed include ggplot2 and dplyr.

## Acknowledgements

We thank Mandy Burns and Amanda Jones for administrative support, Moya Costello for artwork, the Histopathology Shared Resource for pathology support, the Massively Parallel Sequencing Shared Resource and Integrated Genomics Shared Resource for genomics support, and the Oregon Translational Research and Development Institute (OTRADI) for high throughput drug screening support.

## Author contributions

HWL, DSD, SR, AVG, JP, CHX, and SL performed the experiments and analyzed the data. DSD and LMH performed the computational analyses. LC, RW-III, and JMD performed and analyzed the phosphoproteomics data. AR, AVG, SR, and DG analyzed the HTS data. HWL, DSD, CRW, JMD, AR, LMH, and GVT wrote the paper. GVT conceived the project. LMH and GVT supervised all aspects of the work.

## Supplementary Figures

**Supplementary Figure 1:**
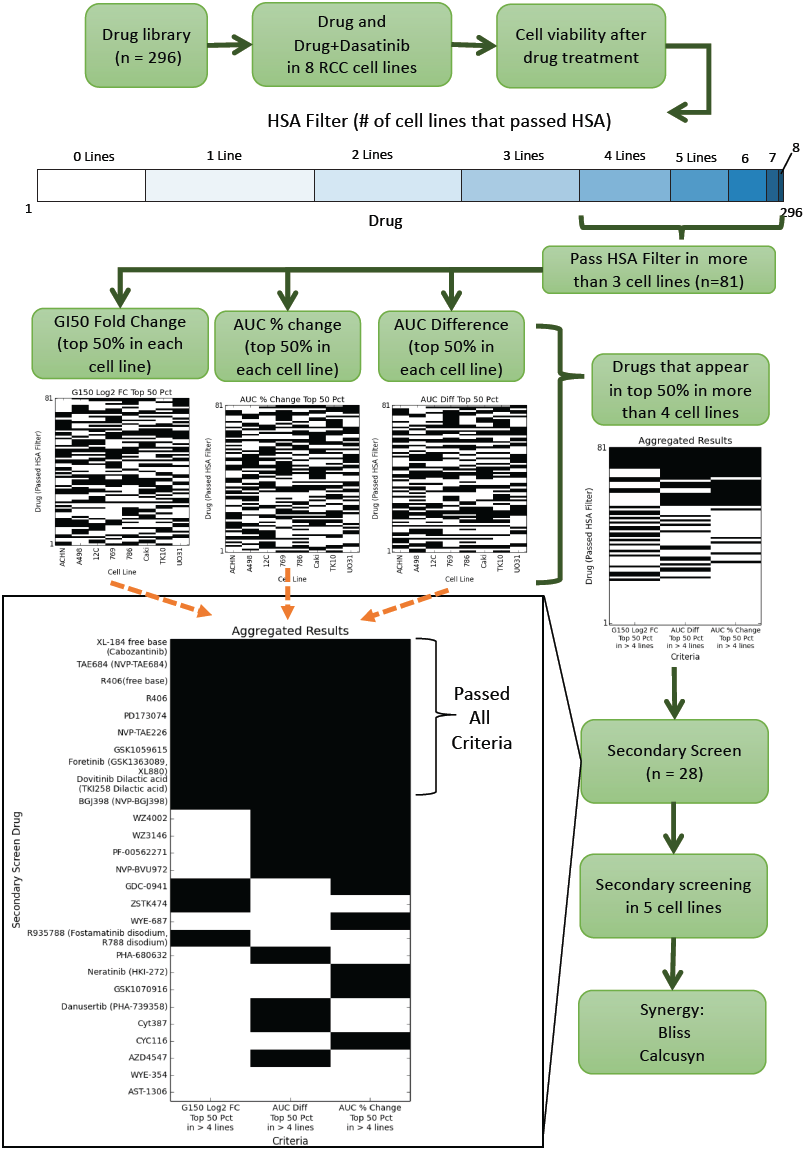
Overview of pipeline for drug selection. The library of 296 drugs were screened in eight cell lines and retained cell viability. Eighty-one drugs passed the “Highest Single Agent” (HSA) filter in four or more lines (the HSA test requires that the combination have at least 10% greater inhibition than either dasatinib or the drug alone at the same dose, for at least three doses). Three measurements of these drugs were calculated for each cell line (G150 fold change, AUC percent change between drug alone and drug+dasatinib, and AUC difference between drug alone and drug+dasatinib). Each measurement results in a matrix where every (drug, cell line) pair is one if the measurement appears in the top 50% of measurements in the cell line and zero otherwise (three inset heatmaps). We then collapse each measurement matrix into a single column, where a drug’s entry is one if more than half of the cell lines have the drug’s measurement in the top 50%. These columns form the criteria matrix (*shown here for three criteria, shown for six criteria in Figure 1B)*. The drugs were shortlisted based on this screen as well as other considerations such as clinical utility, which were subjected to secondary screening in five cell lines.

**Supplementary Figure 2:**
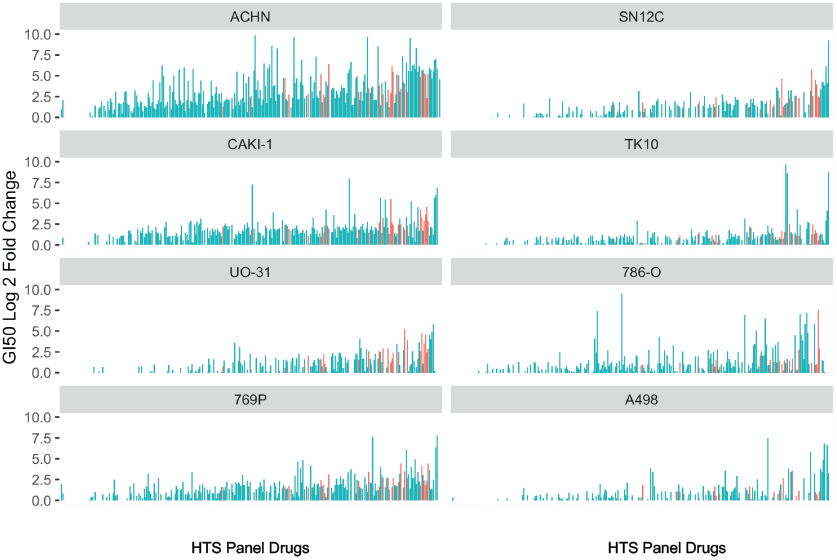
Representation of the fold-change in GI50 for each cell line. Screen data for all compounds tested against human RCC cells: ACHN, SN12C, CAKI-1, TK10, UO-31, 786-0, 769-P and A498. In each graph, the fold-change GI50 that resulted with the addition of dasatinib is represented on the y-axis (capped at 50 for uniformity). Each column on the x-axis represents one drug from the high throughput screen. Those drugs that were selected for the secondary screen are represented in red.

**Supplementary Figure 3:**
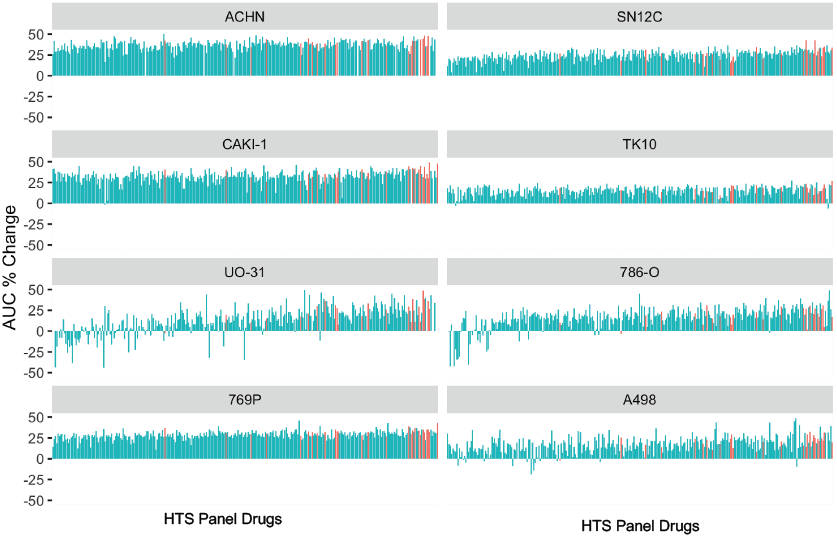
Representation of the percentage change in AUC for each cell line. Screen data for all compounds tested against human RCC cells: ACHN, SN12C, CAKI-1, TK10, UO-31, 786-0, 769-P and A498. In each graph, the % change in AUC (area under dose response curve) that resulted with the addition of dasatinib is represented on the y-axis (capped at 50 for uniformity). Each column on the x-axis represents one drug from the high throughput screen Those drugs that were selected for the secondary screen are represented in red.

**Supplementary Figure 4:**
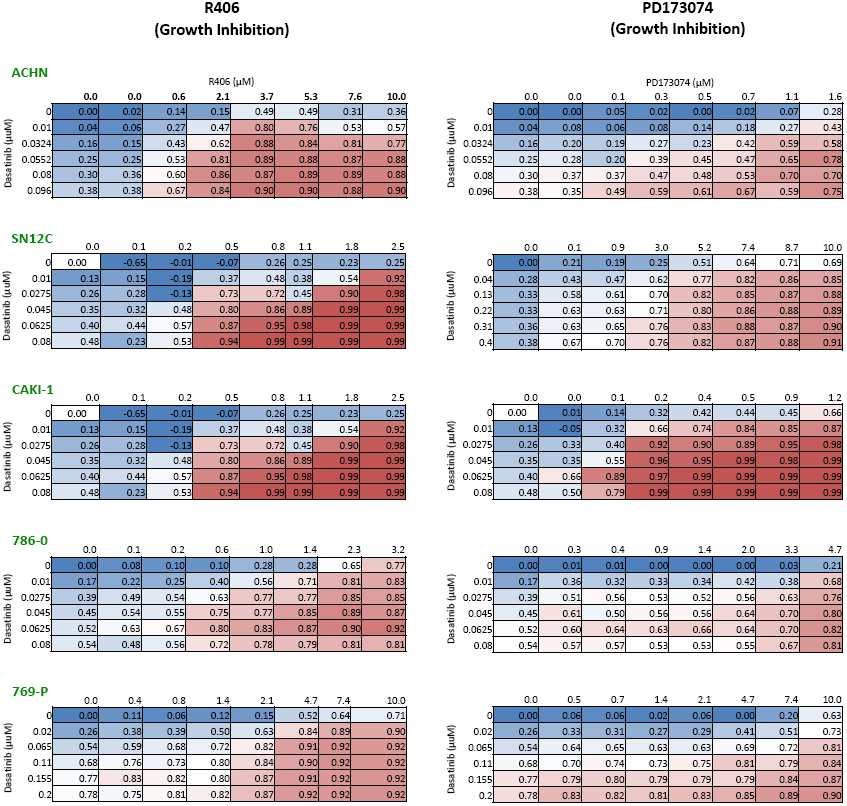
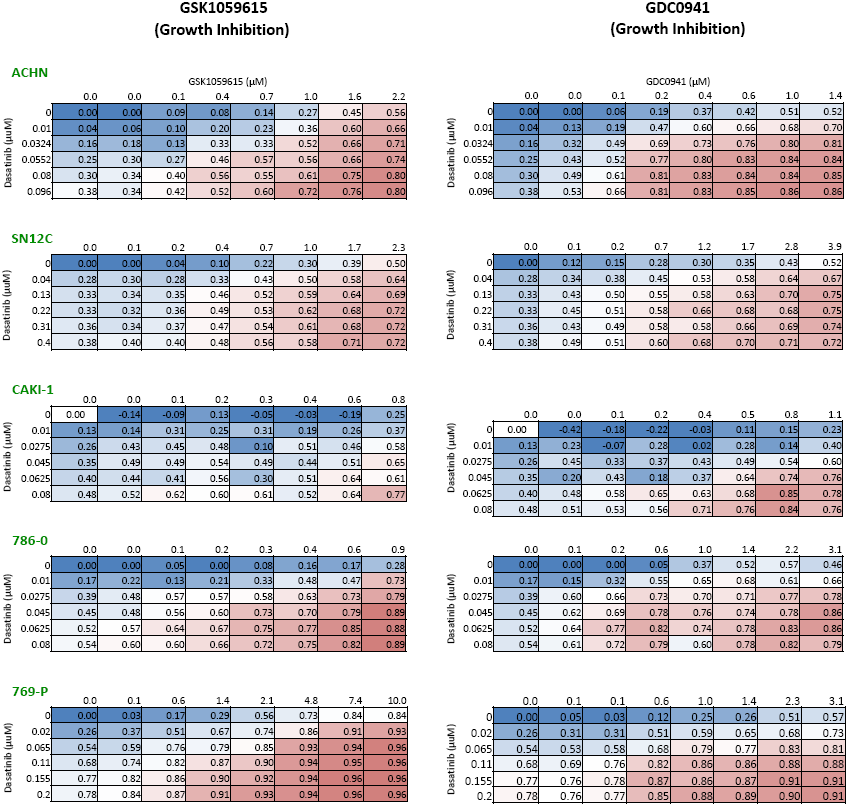
Representation of secondary screening dose matrix. Secondary screening dose matrix of the top ranked synergistic combinations with dasatinib (Calcusyn): R406, PD173074, GSK 1059615, GDC0941 in human RCC cells: ACHN, SN12C; CAKI-1, 786-0 and 769-P. Growth inhibition was assessed after 4 days to varying doses of the stated drugs and dasatinib. Percent inhibition at each dose of the drug is presented.

**Supplementary Figure 5:**
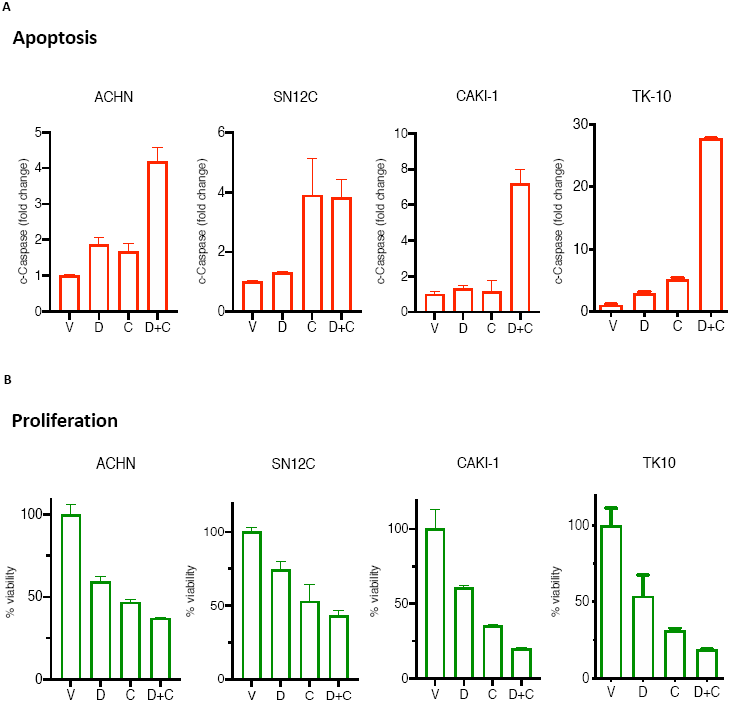
Combination of dasatinib and cabozantinib induces apoptosis and inhibits proliferation. **A.** Cell apoptosis measured by Caspase 3/7 compared to DMSO vehicle control (V) after 72hr-exposure of human RCC cells (ACHN, SN12C, CAKI-1, TK-10) to dasatinib (D), cabozantinib (C) or the combination (D+C). **B.** Cell viability measured by CellTiter Blue compared to DMSO vehicle control (V) after 72hr-exposure of human RCC cells (ACHN, SN12C, CAKI-1, TK-10) to dasatinib (D), cabozantinib (C) or the combination (D+C).

**Supplementary Figure 6:**
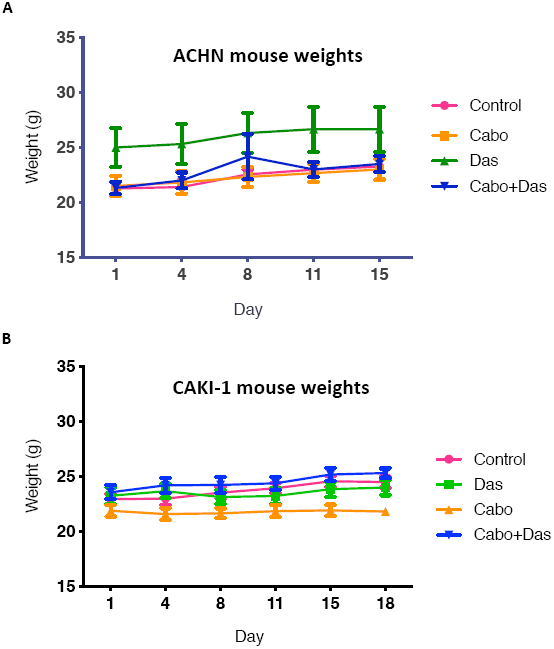
Tolerability of the dasatinib, cabozantinib and the combination in mouse xenograft models. Body weights of mice bearing **(A)** ACHN, and, **(B)** CAKI-1 tumors as indicated. Data are presented as mean ± SEM; *ns*: not significant; ACHN: control vs cabozantinib+dasatinib p=0.4778; CAKI-1: control vs cabozantinib+dasatinib, p=0.8833.

**Supplementary Figure 7:**
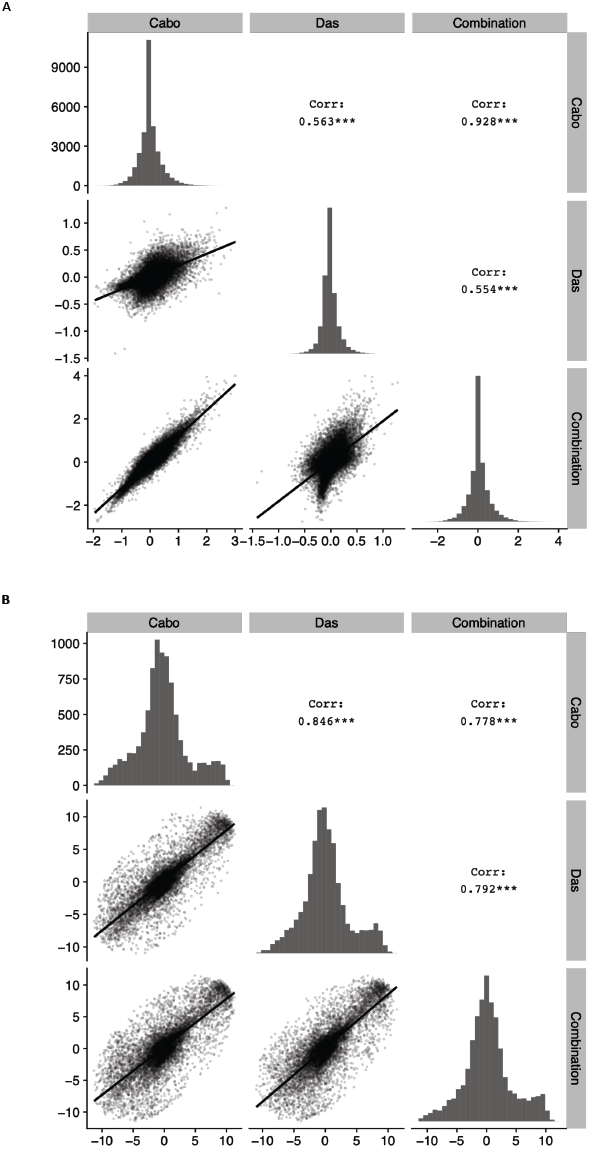
Scatterplots of gene expression and phosphoproteomics. Scatterplots of treatment-induced gene expression changes, in log_2_ scale **(A)**, and phosphoproteomic changes **(B)**, as t statistics, with Pearson correlation R^2^ values, for all pairwise combinations of treatments.

**Supplementary Figure 8:**
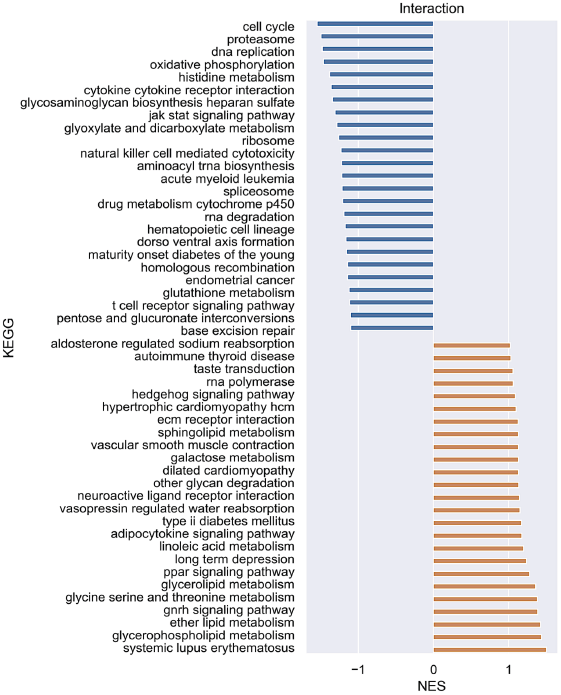
The combination of dasatinib and trametinib (MEK inhibitor) effectively inhibits proliferation in kidney cancer cells. The change in GI50 (on y-axis) that resulted with the addition of dasatinib to trametinib to human RCC cells. The bars are color coded: blue (trametinib) and red (trimetinib+dasatinib).

**Supplementary Figure 9:**
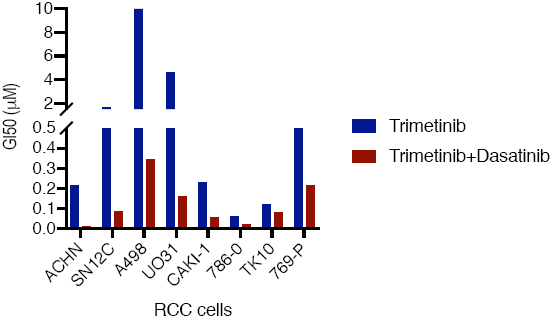
GSEA analysis of the inferred transcriptional master regulators. Gene set enrichment analysis (GSEA) was run on this ranked gene list (Kegg gene set). Normalized enrichment scores were used to generate the bar graph. Top 25 downregulated (blue) and upregulated (red) gene sets from the cabozantinib-dasatinib interactome are shown.

## Supplementary Tables

**Table S1.** Listing of screen drugs

**Table S2.** Dasatinib Vs Control_KSEA_pY

**Table S3.** Dasatinib Vs Control_KSEA_pST

**Table S4.** Cabozantinib Vs Control_KSEA_pY

**Table S5.** Cabozantinib Vs Control_KSEA_pST

**Table S6.** Dasatinib+Cabozantinib Vs Control_KSEA_pY

**Table S7.** Dasatinib+Cabozantinib Vs Control_KSEA_pST

